# Prefrontal control of actions in freely moving macaques

**DOI:** 10.1101/2022.10.26.513892

**Authors:** Benjamin Voloh, David Maisson, Roberto Lopez Cervera, Indirah Conover, Mrunal Zambre, Benjamin Hayden, Jan Zimmermann

## Abstract

Our natural behavioral repertoires include complex coordinated actions of characteristic types. To better understand the organization of action and its neural underpinnings, we examined behavior and neural activity in rhesus macaques performing a freely moving foraging task in an open environment. We developed a novel analysis pipeline that can identify meaningful units of behavior, corresponding to recognizable actions such as sitting, walking, jumping, and climbing. On the basis of action transition probabilities, we found that behavior was organized in a modular and hierarchical fashion. We found that, after regressing out many potential confounders, actions are associated with specific patterns of firing in each of six prefrontal brain regions and that, overall, representation of actions is progressively stronger in more dorsal and more caudal prefrontal regions. Conversely, we found that switching between actions resulted in changed firing rates, with more rostral and more ventral regions showing stronger effects. Together, these results establish a link between control of action state and neuronal activity in prefrontal regions in the primate brain.

## INTRODUCTION

Natural behavior is highly complex, but it is not random; it is organized, although we are only beginning to understand how. It is clear that when we move we strategically select from a repertoire of learned stereotyped actions, and combine these actions into larger sets of action patterns (Merel et al 2019; Berman et al., 2016; Marshall et al 2021). Understanding the organization of behavior and its neural basis has long been an important goal in the fields of ethology, psychology, and neuroscience (Krakauer et al., 2017; Tinbergen, 1951; Gallistel, 2013; Anderson and Perona, 2014; Calhoun and El Hady, 2021; Periera et al., 2020). A greater understanding of behavior would have additional benefits, especially as far as understanding how the brain controls various states and entry and exits into specific action states, which may have psychiatric relevance.

Our understanding of the neural basis of action control is, for the most part, focused on a few key effectors, especially the arm and oculomotor system (Leigh and Zee, 2015; Liversedge et al., 2011; Shenoy et al., 2013; Gallego et al., 2017). Using these methods, classic electrophysiological experiments have identified important roles for primary (M1) and dorsal premotor (PMd) cortices, as well as roles for supplementary motor area (SMA) and even dorsal anterior cingulate cortex (dACC) in action planning and execution (Gallego et al., 2022; Ebbeson and Brecht, 2017; Shenoy et al., 2013; Heilbronner and Hayden, 2016). Other evidence links the ventrolateral prefrontal cortex (vlPFC) with motor inhibition, reflexive orienting, and motor adjustment (Levy and Wagner 2011; Sakagami and Pan, 2007). Finally, some accounts suggest that OFC maintains a representation of extended sequences of actions that aid in fulfillment of a goal (Young and Shapiro 2011). More broadly, some work links the entire prefrontal cortex to control of action; some of this work indicates that there is a ventral to dorsal gradient along which plans are converted to action commands (Fuster, 2000 and 2001; Yoo and Hayden, 2018; Fine and Hayden, 2022; Merel et al., 2019).

However, while a great amount of neuroscience has been devoted to understanding the neural foundations of behavior, nearly all of it has been limited to the study of simple and restricted behaviors, in particular saccades and arm reaches. The study of such behaviors has been extremely important - it has led, among many other things, to great advances in prosthetics, and, by extension, to novel ideas for understanding of cognition. This work has introduced important dynamical systems ideas into neuroscience, which in turn led to advances in other domains (Shenoy et al., 2013; Ebitz and Hayden, 2021; Urai et al., 2022). However, while such behaviors can shed important light onto the neural basis of behavior, they miss the opportunity to tell us about how complex coordinated behavior is mediated by brain activity.

Some recent work has extended neuroscience into the domain of complex behavior (Calhoun and Murthy, 2017; Brown and De Bivort, 2018; Gomez-Martin and Ghazanfar, 2019). For example, recent work shows that behavior of mice is organized into a grammatical structure, and that the dorsolateral striatum mediates this organization (Markowitz et al., 2018). Likewise, selection and change of behavioral state in flies and zebrafish is mediated by specific patterns of neural activity in specific neurons (Calhoun et al., 2019; Marques et al., 2020). Critically in all these studies, and others like them, the ability to measure and identify behavior was essential for obtaining a fuller understanding of neuronal circuitry coordination between regions underlying action control. For this reason, we were motivated to develop a system that could monitor the behavior of freely moving macaques, and therefore allow us to understand how the prefrontal cortex - a region of special interest for decision-making and control - mediates selection of behavior.

The work understanding the organization of behavior has depended critically on the development of high-quality video-based motion tracking systems. These systems have led to the ability to track positions of body landmarks in small animals, including worms, flies, and mice (Mathis and Mathis, 2020; Periera et al., 2020; Calhoun et al., 2019; Sturman et al., 2020; Hsu and Yttri, 2020; Bohnslav et al., 2021; Calhoun and Murthy, 2017). This problem is much more difficult in primates, although, even here, significant progress has been made (Marks et al, 2021; Dunn et al., 2021; Bain et al., 2021; Bala et al., 2020; Labuguen et al;, 2021; reviewed in Hayden et al., 2022). In smaller animals, the behaviors identified by tracking systems allow for the automated identification of specific meaningful behavioral units (sometimes called “ethogramming”, Marshall et al., 2021; Berman et al., 2016; Wiltschko et al., 2015; Bain et al., 2021; Bohnslav et al., 2021). Behavior in these organisms consists of structured sets of behaviors that are organized hierarchically (Berman et al., 2016; Marshall et al., 2021; Wiltschko et al., 2015). The success of these methods raises hopes that we can develop parallel methods for quantifying behavior in larger animals with more complex repertoires, such as macaques.

However, doing so requires novel analytical approaches because macaques’ bodies have much higher degrees of freedom (Bala et al., 2020). Despite this difficulty, macaques are especially important in this regard because of their pivotal role as a model organism for biomedical research (Rudebeck et al., 2019; Buffalo et al., 2019).

We examined the behavior of two macaques performing a foraging task moving around a large (2.45 × 2.45 × 2.75 m) open field. We made use of OpenMonkeyStudio, a system that can perform detailed three-dimensional behavioral tracking in rhesus macaques with high spatial and temporal precision (Bala et al., 2020). We used this system to track the position of 13 joints at high temporal and spatial resolution as our subjects performed three different behavioral tasks. We recorded brain activity in six brain regions, orbitofrontal cortex (OFC), dorsal anterior cingulate cortex (dACC), supplementary motor area (SMA), ventrolateral prefrontal cortex (vlPFC), dorsolateral prefrontal cortex (dlPFC), and dorsal premotor cortex (PMd). In all regions, we found that neural activity varies systematically with the state of the subject. We found that the strength of state coding was systematically greater in more caudal and more dorsal structures. We also found that neurons have non-specific signals associated with the switch between actions, and that the strength of these signals is - in a reverse of the pattern for action signals - progressively stronger in more ventral prefrontal structures.

## RESULTS

### Behavioral and neural recordings

We studied the behavior of two rhesus macaques (*Macaca mulatta*) performing a depleting-rewards *freely-moving foraging task* (see **Methods**) in a large open cage (2.45 × 2.45 × 2.75 m cage with four barrels) that allowed for free movement **(Figure 1A and B**). The enclosure contained reward stations (typically four, but on a few occasions two or three; see **Methods**). These reward stations consisted of touchscreens, juice reservoirs, dispenser tubes, and levers (**Figure 1B**). Each station had the same arrangement, but individual stations were positioned at fixed locations of varying heights. Subject behavior was otherwise physically unconstrained within this large environment.

**Figure 1.**
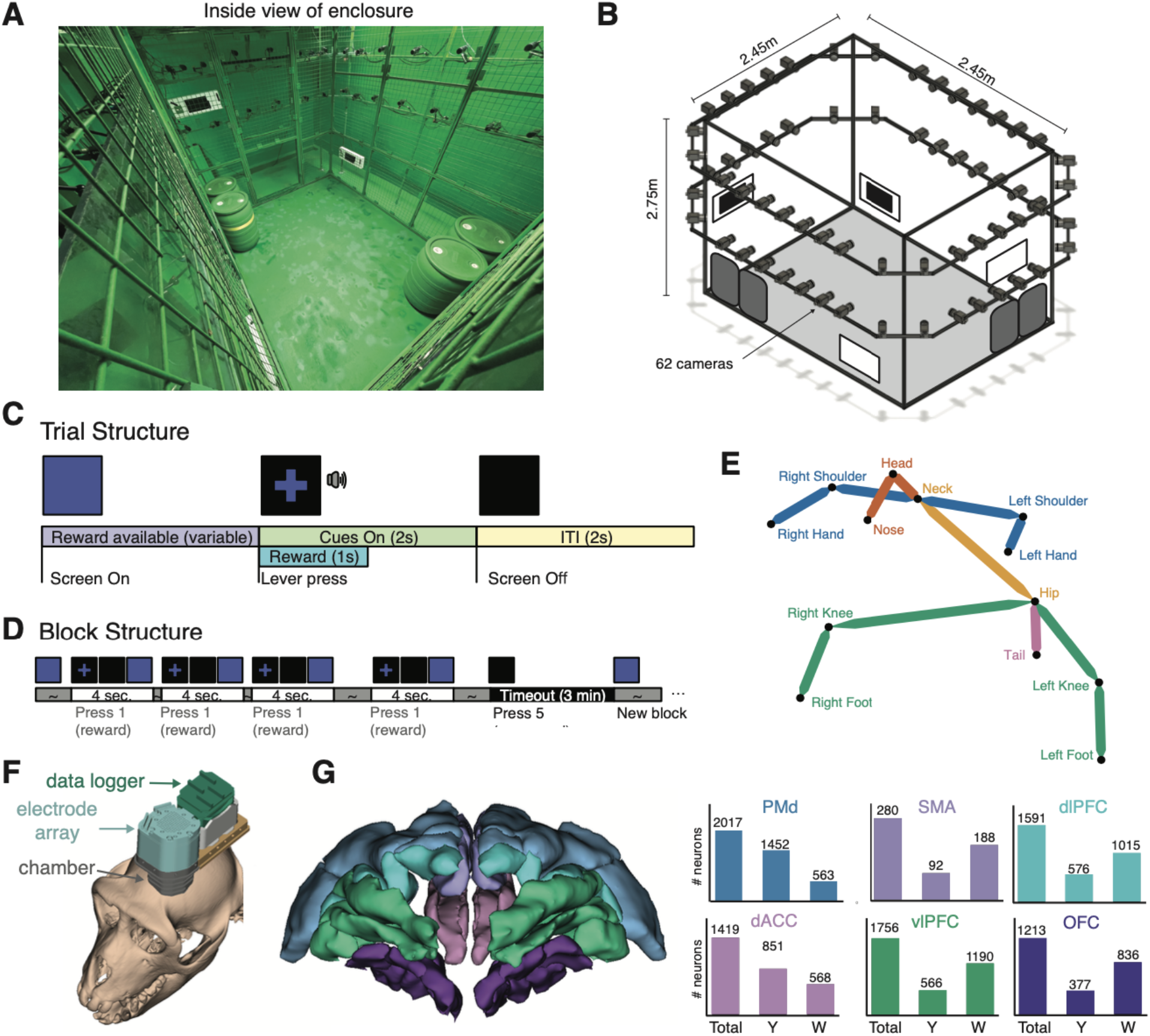
Environment, task, and electrophysiology for freely moving macaques. **(A)** Photograph of inside of enclosure permitting freely moving foraging. Up to four feeders were present for the experiment, located at different corners and requiring different postures to reach. **(B)** Schematic of the cage. 62 machine vision cameras provided multi-view coverage of every part of the cage. **(C)** Structure and timing of one trial. Following a blue initialization screen, subjects pressed a lever and received fluid reward. After two seconds, the trial reset **(D)** structure of one block of trials. Four rewards were available. A fifth lever press initiated a 3-minute timeout period. **(E)** 3D reconstruction of 13 joints using the OMS pipeline. **(F)** 3D model of the recording system superimposed on a subject’s cranium. **(G)** 3D rendering of the prefrontal areas from which neural data were recorded (left), and the corresponding number of recorded neurons (right).

The task was straightforward; the computer displays were initially all blue. If the subject pressed the lever at a reward station, the juice tube provided an immediate aliquot of preferred liquid reward (typically water, always 1.5 mL) and a green cross appeared while a two-second tone played (**Figure 1C**). When the tone stopped, the screen turned off. Following another two-second interval, the screen returned to blue to indicate that the lever had been reactivated. The subject could repeat this process four times to obtain four (identically sized) rewards (**Figure 1D**). In order to replenish rewards, a fifth lever press was required, which deactivated the reward station for three minutes.

The length of each daily recording session was determined by the subject, and typically lasted 100-120 minutes (average 100.2 minutes). We analyzed 81 sessions in subject Y and 86 in subject W (*see* **Methods** for exclusion criteria). Subjects were tracked with 62 high-resolution machine vision cameras (**Figure 1B)** and the pose (3D position of 13 body landmarks, see **Methods**) was determined for each frame (30 frames per second) using our pose-tracking system, *OpenMonkeyStudio* **(Figure 1E;** Bala et al. 2020). This system provides estimates of the positions of 13 body landmarks in every frame of video in three dimensions. As a result, we obtained an average of 180,307 frames of pose per session, and a total of 35,881,005 frames. We simultaneously recorded neural activity using a locally mounted data logger attached to a multi-electrode (n=128 electrodes, **Figure 1F**) array with independently moveable electrodes (**Figure 1F, Methods**), targeting multiple regions in the prefrontal cortex (**Figure 1G)**

### Embedding recovers actions

To identify behaviors, we used an embedding approach similar to one that has been used successfully in rodents and flies (Berman et al 2014; Marshall et al 2021). A high-level description of the pipeline is visualized in **Figure 2A**. In brief, we extracted kinematic features from the thirteen-joint reconstructed pose, reprojected these samples onto a common two-dimensional embedding space, and clustered samples to discover repeated, recognizable behaviors. We extracted a total of ninety-one various features in both an egocentric and allocentric reference frame. These included limb and joint configurations, periodicity in limb configurations, and gross dynamics of the subject; *see* **Methods** for a full list of features, and **Figure 2E** for an example).

**Figure 2.**
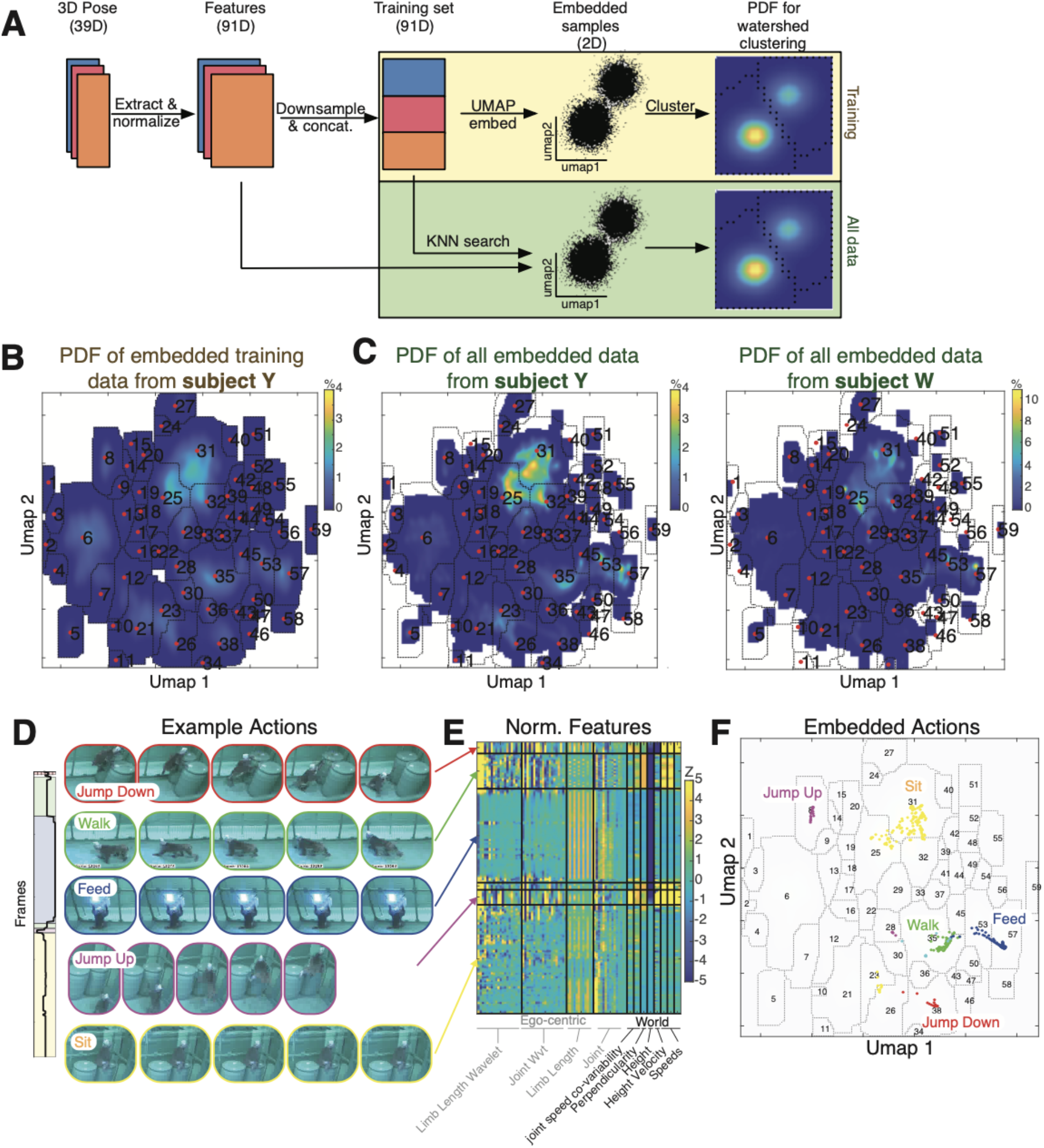
Embedding recovers actions. **(A)** Pipeline for unsupervised behavioral segmentation. After preprocessing, each dataset of 13 joint positions was used to extract 91 salient features. We generated a 2D embedding, which was used for clustering via watershed. Using the learned embedding and cluster labels, we used the training set to re-embed and re-label all data. **(B)** Probability density map of Subject Y. Peaks in the heatmap: samples with similar postural dynamics. Dotted lines: cluster boundaries from the *watershed algorithm*. Numbers: cluster labels. **(C)** Same as (B), but for the (re-embedded) testing data for subjects Y (left) and W (right). **(D)** Example actions from one dataset with an ethogram of actions. (left) ethogram of actions. Colored backgrounds represent different actions as determined by an experimenter. Black line represents the learned cluster label. **(E)** Normalized features computed from pose kinematics from the examples in (D). Color limits have been scaled to aid in visualization between features. Features were computed in both ego-centric and world reference frames. **(F)** Embedding density map from (B), with data from the example actions (D,E) overlaid. Actions were well separated from each other.

To cluster samples robustly and efficiently, we employed non-linear dimensionality reduction (UMAP; McInnes et al 2018) on these 91 features to reproject samples. We trained the embedder on a training set from subject Y. We chose to construct our basis set from the subject with greater behavioral range; had we chosen the other, we would have obtained similar, albeit slightly noisier, results; data not shown). This process resulted in a reprojection of the 91-dimensional behavioral space onto two dimensions, in which periods of similar kinematics are adjacent to each other in the resultant lower dimensional space.

We then performed a kernel density estimation to approximate the probability density of embedded poses at equally interspersed points (**Figure 2B**). The embedding density maps clearly reveal clustered organization, as evinced by visually separable peaks (**Figure 2B)**. Each cluster reflects a set of similar poses that are relatively distinct from others. We formally identified clusters using a *watershed algorithm* on the density map from the training set (technically, on its inverse, so that peaks became troughs; Berman et al. 2014). This algorithm treats troughs in the maps as sinks and draws optimal boundaries separating them. The resulting embedding space contained fifty-nine distinct clusters. We then used the learned embedding and cluster assignment to reproject all data samples from both subjects and label each action accordingly (**Figure 2C**).

Examples of behaviors discovered through this pipeline are visualized in **Figures 2E-F. Figure 2D** shows example behaviors delineated by an observer (such as jumping, walking, and sitting), and the corresponding cluster assignment. Each of these behaviors was associated with a unique fingerprint of kinematics (**Figure 2E)**. As such, these periods corresponded to well-separated clusters in the low-level embedded space (**Figure 2F**). This provides confirmation that the extracted clusters corresponded to recognizable and nameable actions. As such, for the remainder of this paper, we will refer to these clusters of behavior as *actions*. Each action lasted on average 1.63+0.009 sec. in subject Y, and 0.67+0.0025 sec. in subject W.

### Action organization is modular and hierarchical

It has long been supposed that natural behavior in primates has an inherently hierarchical organization. We next sought to characterize this hierarchy in our subjects. Formally speaking, we asked if *sets of actions* tend to co-occur, so that just as the co-occurrence of poses produces actions, the co-occurrence of actions would produce higher level organization. To do this, we computed transition probability matrices for each possible pair of actions using the actions discovered through embedding (see above).

The transition probability matrix this process produces is, in graph theoretic terms, a directed graph. In the parlance of graph theory, nodes that form strong links between each other are referred to as modules or communities. Because of this, we refer to sets of actions that have a high probability of co-transition as “*action modules*”. The identification of these modules allows us to sort the transition probability matrix so that the actions that are part of the same module are adjacent; graphically, the result of this is that there will be conspicuous blocks on the diagonal (**Figure 3A**). These blocks correspond to action modules.

**Figure 3.**
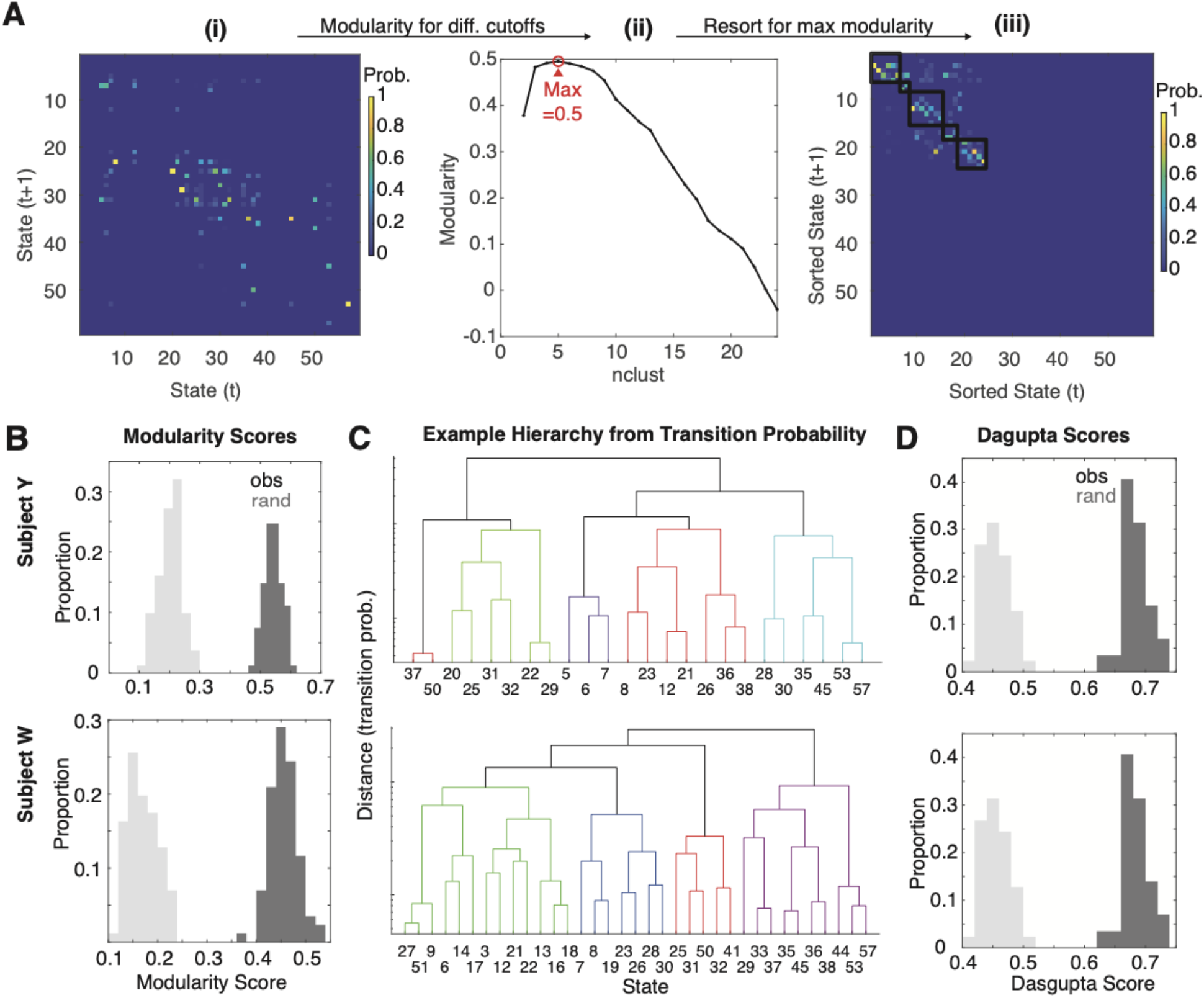
Actions show modular and hierarchical organization. **(A)**. Example modularity for one dataset. **(Ai)** Original transition probability matrix with a transition lag of 1. **(Aii)** Modularity score for different cuts of the behavioral hierarchy. The maximum modularity was for 5 modules. **(Aiii)** Transition matrix sorted to provide maximum modularity. It is evident that transitions tend to occur within, rather than between, modules. **(B)** Distribution of maximum modularity (e.g. peak in Aii) for each dataset, for subject Y (top) and W (bottom). Observed modularity scores (dark) were greater than chance for all datasets (light). **(C)** Example dendrogram showing action organization for one example session for each subject **(D)** *Dasgupta scores* for each dataset, measuring the degree of hierarchy as evidenced by transition probabilities. Observed *Dasgupta scores* were greater for observed (dark) rather than randomized (light) transition matrices, indicative of the existence of a hierarchical structure.

We then used a recently developed algorithm named *Paris* (Bonald et al, 2018), which performs hierarchical clustering on the graph derived from transition probabilities and returns a tree (we will examine the assumption of hierarchy using the Dasgupta score below). To determine the number of modules, we cut the tree at several levels and computed a *modularity score* for the resulting subtrees, and chose the number of modules that maximized the modularity score. The modules that result from this cut give the highest average within-module action-transition probability and the lowest average across-module action-transition probability. These modules maximize the difference between these two measures. From this process we can identify the most likely behavioral modules (that is, the ones that fit the data the best).

An example of this process is visualized in **Figure 3A**. For the transition probability matrix (**Figure 3Ai**), modularity was maximized for 5 modules (**Figure 3Aii)**. The modular nature of transitions is evident in the sorted transition probability matrix (**Figure 3Aiii**). Note that not all actions were present in this session; we performed clustering and modularity computations with this in mind.

We tested for the modular organization of behavior by computing the modularity score for each session. We found that all sessions, in both subjects, exhibited significant modularity (**Figure 3B**; randomization test, p<0.001). In other words, behavior was organized such that specific modules of actions tended to transition between one another, but not actions outside of the module. The average number of modules was 4.6+0.08 in subject Y, and 4.83+0.07 in subject W. These results indicate that subjects’ behavior was organized into modules, each of which consisted of stereotyped actions.

We then asked if action modules exhibit higher level organization. Examples of hierarchically organized actions are shown in **Figure 3C**. Higher level connections in this dendrogram show how different action modules are related. To quantify the degree of hierarchical organization, we computed the *Dasgupta score*, which quantifies the quality of hierarchical clustering on the transition probability graph (Dasgupta 2016). A score above chance indicates the observed tree has high level components that are distinctly related to one another. The *Dasgupta score* was above chance in all sessions in both subjects (**Figure 3D**; randomization test, p<0.001), indicating hierarchical organization of behavior.

### Neurons across the prefrontal cortex encode actions

We next sought to understand how actions are encoded across six prefrontal regions. We recorded a total of 10,502 neurons over 196 sessions. Of these, 2,818 neurons were excluded from the following analyses because preliminary investigations indicated that their sessions had tracking that was too noisy for our purposes, or regressing out extraneous variables failed (see **Methods** for exclusion criteria). The remaining 7,684 neurons were recorded from six structures in the prefrontal cortex, OFC, vlPFC, dACC, SMA, dlPFC, and PMd (*see* **Methods** and **Figure 1E**).

We examined the average firing rate of each neuron during each of the fifty-nine identified actions by averaging each neuron’s responses across the entire action (see **Methods**). To isolate activity related to actions, we fit a Poisson generalized linear model (GLM) to identify neuronal responses related to task events (such as lever presses, rewards, and cues) and subject position (in the XYZ dimensions). We used the fit model to residualize the neuronal time-series, and then downsampled the residualized time-series to match that of the pose (30 Hz). All neural analyses were performed on these residualized, downsampled time-series.

We found that neurons exhibited specific firing to individual actions. An example of rates of neurons recorded in one session (n=79 neurons) to six different actions is shown in **Figure 4A**. In this session, neurons had distinct specific firing patterns for each action, and different areas showed distinct patterns as well (**Figure 4B**). We operationalized the strength of action encoding of each individual neuron by performing a Kruskal-Wallis test and extracting the X^2^ value. This value reflects how well actions can be dissociated from one another by the (residualized) firing rate. We opted for a non-parametric measure as rates were highly non-uniformlly distributed. In this session, posterior regions, such as the PMd, had the highest average strength of action encoding while more ventral and anterior regions, such as dACC, vlPFC, and OFC, had weaker encoding (**Figure 4B**).

**Figure 4.**
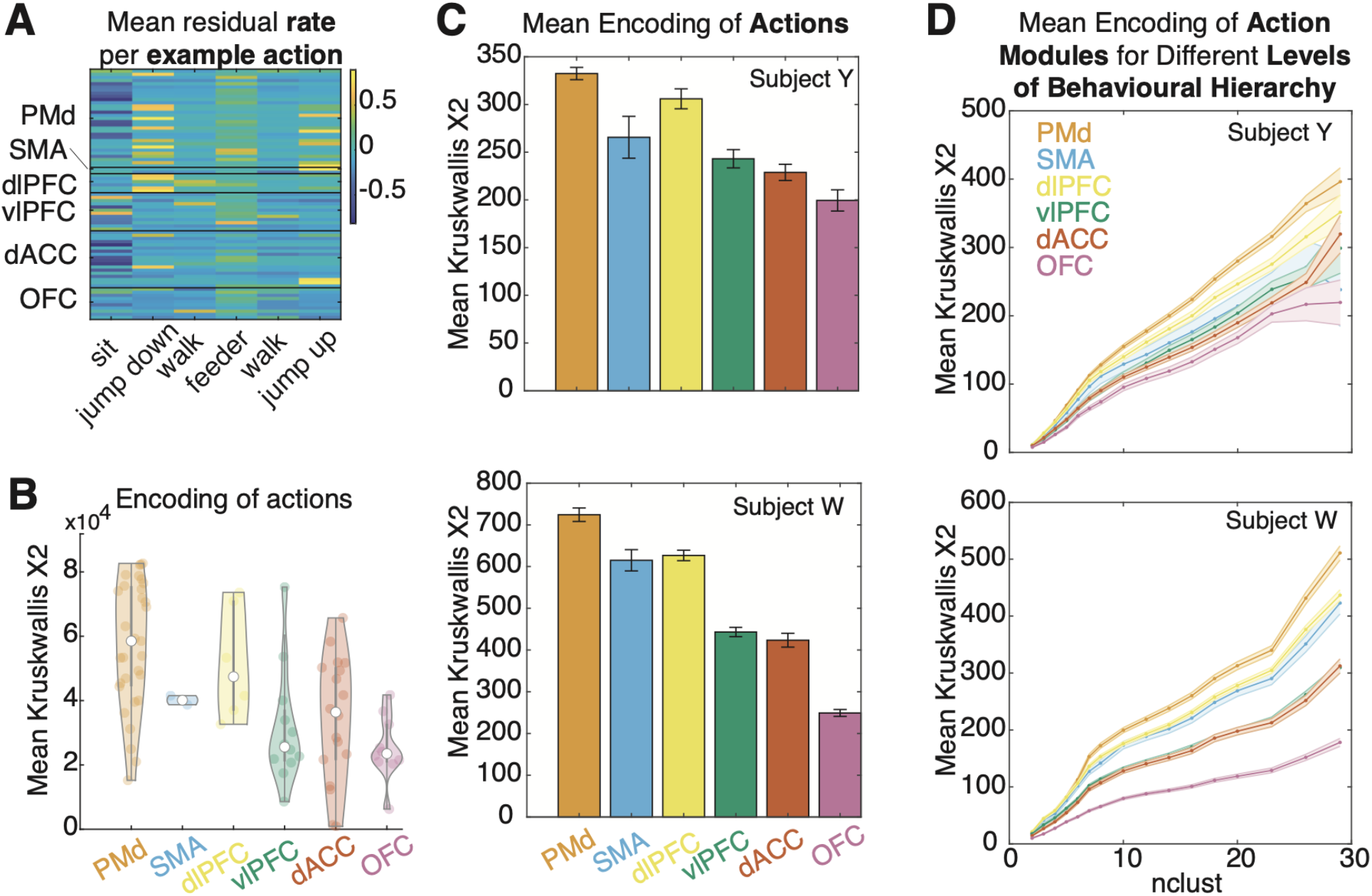
Actions encoding across brain areas. **(A)** Mean (residualized) firing rates for six example actions from one session, split by brain area. Here, residualized means after accounting for potential confounders (see **Methods**) **(B)** Mean encoding strength for each area for the entire session, operationalized as the Kruskal-Wallis chi-squared value, across all recorded areas. More dorsal areas exhibit higher encoding strength. **(C)** Mean encoding strength across all sessions and areas, in subject Y (left) and W (right). More dorsal areas exhibited stronger encoding strength. **(D)** Similar to (C), but for different cuts of the behavioral hierarchy (i.e. behavioral granularity). The same inter-areal effects are present for different levels of behavioral granularity, in both subjects Y (left) and W (right).

Indeed, across all sessions and in both subjects, we found evidence of significant action encoding in all six regions tested (**Figure 4C**). Specifically, in all 6 regions, the median X^2^ was significantly greater than chance (randomization test, p<0.001 in all areas and for both subjects), whereas randomized data showed a chi-square at chance levels). Furthermore, the mean strength of encoding differed between regions (ANOVA; subject Y, F=32.2, p<0.001; subject W, F=171, p<0.001). These results suggest that action encoding is prevalent across the prefrontal expanse, even after controlling for task related activity and subject position.

As noted above, actions are organized into action modules consisting of clusters of action that tend to co-occur. We next asked whether neural activity was selective for action modules. To this end, we remapped cluster labels to encompass larger action modules by cutting the transition probability dendrogram at increasingly coarser action module levels, and re-calculating the encoding strength (**Figure 4D**). We found that even for very coarse descriptions of behavior (even with as few as two action modules), neural activity encoded action modules as well (randomization test, multiple comparison corrected, p<0.001 for all areas, number of clusters, and subjects). In other words, neural activity recapitulates, at least at a very coarse level, the structure of behavior. The inter-areal pattern of the strength of encoding remained the same, with PMd showing the strongest encoding and OFC showing the lowest.

### Action encoding strength grows along a dorsal-ventral gradient

A visual inspection of the data suggests that action coding grows systematically with the dorsoventral position of the recording site (**Figure 4C**). We next sought to formally test this hypothesis by modeling encoding strength as a function of electrode position (**Figure 5A**). For each electrode, we extracted its depth (Z position, along the vertical axis) and its X (along the coronal plane) and Y (along the sagittal plane) position on the recording grid. We then fit a linear model, predicting encoding strength (specifically, X^2^ value) as a function of 3D spatial position (see **Methods**).

**Figure 5.**
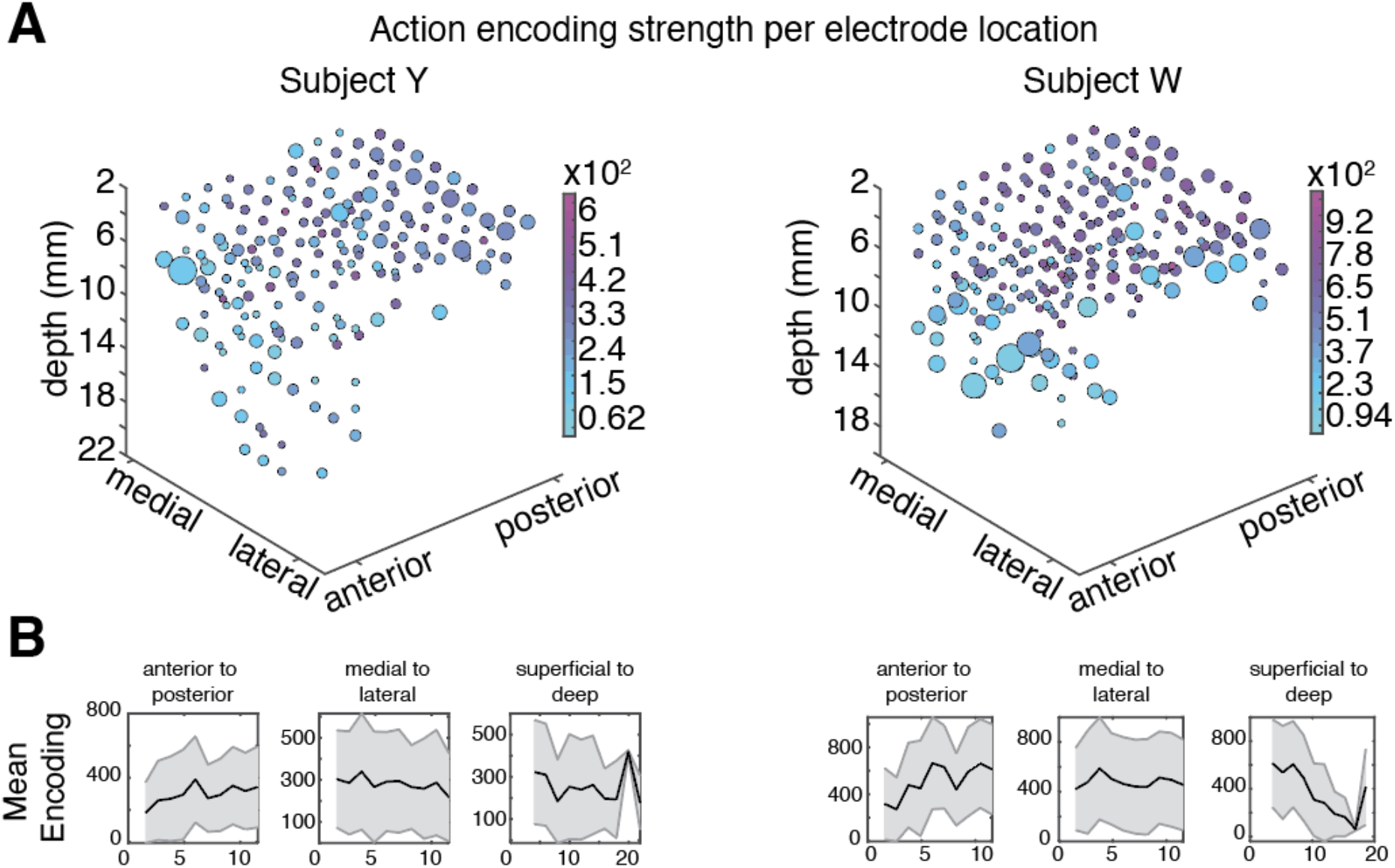
A dorsal-ventral gradient of action encoding. **(A)**. Average encoding strength as a function of electrode position for subject Y (left) and subject W (right). Bubbles correspond to bins of neurons. Color: strength of encoding; size: number of neurons in bin. Stronger encoding (darker color) is more common in dorsal and posterior locations; weaker encoding (lighter color) is more common in ventral and anterior locations. **(B)** Average encoding strength after collapsing along the anterior-posterior (left), medial-lateral (middle), and superficial-deep (right) axes.

We found that encoding strength varied systematically with the position of the electrodes (subject W; F=293, p=0, R2=0.18. Subject Y; F=70, p=0, R2=0.05). Looking specifically at each dimension, we found a significant positive relationship between coding strength and elevation (stronger coding in more dorsal locations; subject W, p<0.0001; subject Y, p=0.02). We also found a significant positive relationship between encoding strength and anterior-posterior position (stronger coding in more caudal locations; subject W, p<0.0001; subject Y, p=0.0003), We did not find any relationship along the mediolateral axis in either subject (subject W, p=0.28; subject Y, p=0.52, **Figure 5B)**.

Some locations had much better coverage than others, which could lead to spurious fits due to large imbalances in the data. To control for this possibility, we binned encoding values according to spatial location (for 5, 10, 15 and 20 bins along each dimension), and then repeated this analysis. We found the same pattern in all cases (data not shown). We conclude that, at least for our analysis methods, actions are most strongly encoded in posterior and superficial areas, such as PMd, and weakest in deep anterior structures, such as OFC.

### Relative increase in firing rate grows for more ventral structures

We next wondered how neurons implement switches between actions - do they simply identify the action being implemented, or do they signal the switch as well? We therefore next examined the patterns of neural activity associated with switches between actions and action modules.

To this end, we selected a two-second time window centered on the moment of switching around each action transition. Since for this analysis we considered peri-switch activity, for any one neuron, we only considered segments where the action before or after the switch lasted longer than 200 ms, thus ensuring that the 400 ms around the switch were not contaminated by other switches. We then normalized activity within and across neurons (see **Methods**). We then assessed the effect of switching on firing, by z-score normalizing each peri-switch segment. We calculated the median and median absolute deviation in the pre-switch period and used these to normalize the entirety of the segment (we opted for these robust measure of central tendency and spread because the data were highly skewed). Then, by taking the median across all segments, we extracted the segment-normalized activity around action switches for each individual neuron.

The average segment-normalized activity is shown in **Figure 6**. We found that in all six cortical regions, and in both subjects individually, neuronal activity changed after action switches. In OFC, which saw the strongest switch effects, the switch produced a reliable increase in neural activity in both subjects (0.02-0.06 times greater than the pre-switch mean; **Figure 6B**). The type of change differed somewhat between subjects. PMd, SMA, and dlPFC were inconsistent between subjects, showing relatively increased activity in subject Y, and decreased activity in subject W. On the other hand, PMd, vlPFC, and dACC showed a similar trend to increase activity in both subjects. However, in both subjects, the average difference between pre and post-switch activity differed by area (Kruskal Wallis test; subject Y, X2=48, p<0.001; subject W, X2=176, p<0.001). Furthermore, the difference in activity could be predicted by the position of the electrodes (linear model; subject Y, F=10.3, p<0.001; subject W, F=49.1, p<0.001). The difference was greater for deeper depths (subject Y, p<0.001; subject W, p<0.001), and for more anterior (near significant in subject Y, p=0.06; subject W, p<0.001) locations. In subject W, the difference was also greater for more lateral locations (p<0.001); this effect was not significant in subject Y, though the effect grew along the same direction (p=0.2). Thus, the relative change in activity after action switches exhibited the opposite trend as for action encoding *per se*, with more ventral regions showing greater activity post-switch, and more dorsal regions showing lesser or relatively lesser activity post-switch.

**Figure 6.**
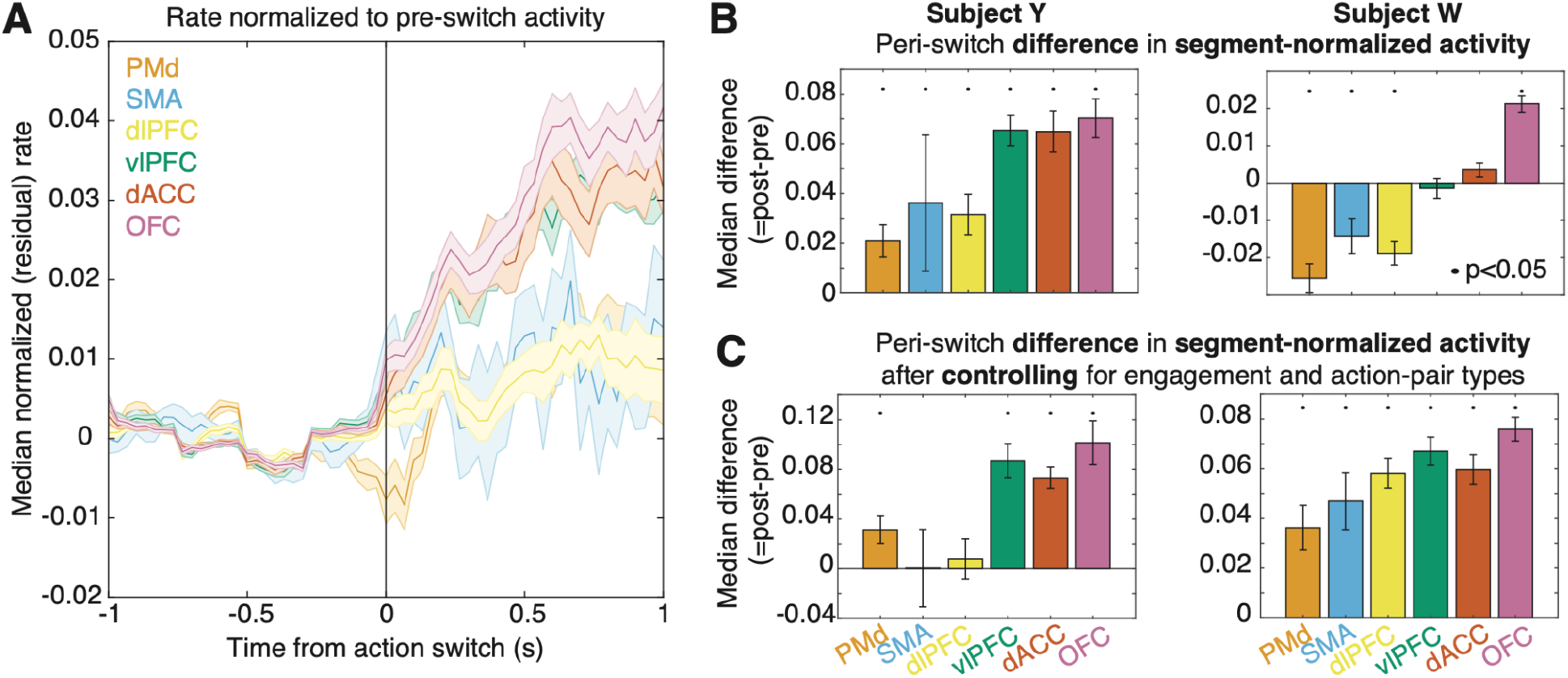
Neural correlates of action switches. **(A)** Average segment-normalized activity aligned to time of switch. Colors indicate brain areas. **(B)** Average (median) difference of post vs pre segment-normalized activity, split by area (color). Black dots: significant difference (p<0.05). Both subjects show an increase with more ventral structures **(C)** Same as (B) but after controlling for possible reward and action-pair type effects.

We considered multiple potential confounds to this analysis. First, we may have failed to fully regress the effects of task and reward. More ventral regions such as OFC and dACC have been particularly strongly associated with reward processing in the literature (Wallis, 2007; Padoa-Schioppa, 2007; Heilbronner and Hayden, 2016). Thus, the relatively higher activity evident after action switches may have been due to engagement with the task. Second, there is a large imbalance in possible action transitions (*see* **Figure 3**). Although this is why we opted for the segment-normalized activity, large imbalances may nevertheless affect the result. To control for these possibilities, we did two things. First, we only considered action transitions in periods of time when subjects were not engaged with the task for at least 1 second. Second, we calculated a weighted average of activity, whereby we first calculated *action-specific* peri-switch activity for each possible pre-switch action, and then averaged across these. After performing these controls, we found a similar pattern of results; namely, that there was a greater increase in post-switch activity in more ventral regions such as OFC, and a more modest (or non-existent) change in more dorsal/posterior regions such as PMd(**Figure 6C**). We also found that the directionality of results was more consistent between subjects, particularly for PMd, SMA, and dlPFC.

## DISCUSSION

Here we investigate the behavior and neural correlates of behavior in two rhesus macaques performing a foraging task in a freely moving context. Our analysis pipeline used a set of kinematic features calculated over short time windows and a dimensionality reduction approach to clustering to identify fifty-nine distinct actions. On the basis of transition probabilities between actions, we were then able to determine that actions were organized into action modules, and to further delineate the full hierarchical structure of this space of actions.

We showed that responses of neurons in six brain regions encoded actions, with greater encoding in more dorsal structures. We found that neurons show a characteristic signature of action-switching (a rise in overall activity), and that the magnitude of this signal is greater in more ventral structures.

Perhaps the most striking finding is the two oppositely pointed hierarchies in prefrontal cortex. Several observers have proposed that the prefrontal cortex is characterized by a ventral-to-dorsal functional gradient in which information is transformed from abstract to concrete, and from conceptual to motor. Partially supporting this idea, we have shown stronger task (Maisson et al., 2021), navigational (Maisson et al., 2022), and action-specific (this paper) representations in more dorsal structures. However, we have not previously identified any variables that have stronger patterns of modulation in more ventral structures. We do that here. Specifically, we show that more ventral regions have a stronger task-switch signal. This signal appears to start before or around the time of the switch, and is common across different switch types, suggesting it is either amodal or has an amodal component. This result, then, is consistent with other proposed task-switching signals, such as those observed in prefrontal and parietal circuits (Dove et al., 2000; Sohn et al., 2000; Nakahara et al., 2002; Rushworth et al., 2002, Brass et al., 2005; Crone et al., 2006; Hyafil et al., 2009). One explanation for these results may be that the switch reflects the coordination of activity across multiple regions, but with the bulk of the relevant cognitive processing most ventrally; another would be that the switch signals are more latent in more dorsal structures, perhaps because the switch signals regulate ongoing hierarchically intermediate neural activity.

Some recent work has begun to unravel the neural processes that occur during free movement (Calhoun et al., 2019; Markowitz et al., 2019; Marques et al., 2019). Our work builds on this past work in several ways. First, we are able to examine more complex behavior that involved more degrees of freedom and more complex planning; as such, we were able to measure neural correlates of actions in an animal with a richer behavioral repertoire (59 distinct actions). Second, we are able to draw conclusions about the organization of the primate prefrontal cortex, whose homology to other species remains debated (Laubach et al., 2018; Heilbronner et al., 2016; Passingham and Wise, 2012). Third, and relatedly, because of our ability to record in six different regions across most of the prefrontal cortex, we were able to link encoding of state and state switching to a specific prefrontal hierarchy (Fuster, 2020 and 2021).

One limitation of this work is that, while we find neural correlates of action, we do not investigate the specific type of information carried in these signals. It may be that neural activity specifies the details of action in a very detailed way, such as describing where each limb should be and what joint angle should be generated (cf. Fetz, 1992). Conversely, it may be very abstract -it may encode the action, or even entire sequences of action, and let downstream structures specify the details. For example, in a study of *C. elegans*, Marques and colleagues found that sensorimotor tuning accuracy changed markedly with the animals state (Marques et al., 2019). These are critical questions that should be addressed in further study of unrestrained and spontaneous behavior (ideally in a setup such as in the present study), but are distinct from the points of interest of the present work. Specifically, we did not seek to tackle how actions are planned, executed, and monitored, but rather sought to obtain a heuristic comparison of action encoding across the brain. By delineating that actions are more strongly encoded in dorsal structures, whereas more ventral ones have a stronger signal for action switching, this work suggests that future studies may focus on ventral structures for action planning and monitoring, and dorsal ones for execution.

Recently, several studies of simpler animals have established the “physics of behavior”, namely, the core elements from which complex behavior is composed, and the rules that govern how those elements combine (Brown and De Bivort, 2018). So far, this approach has not been applied to non-human primates; instead, study of the behavior of non-human primates has largely been limited to highly constrained and simple motoric experiments (such as reaching and saccades), or otherwise rely on simple heuristics to delineate gross motivational state (Shahidi et al 2021). Our study builds on this work to discover specific actions via an automatic and unsupervised approach and delineates their organizational principles. While it is not surprising that behavior is hierarchically organized, it is important to have a specific map of the important behaviors.

One of the greatest potential benefits for statistical analysis of highly quantified behavior is in the prospect of automated ethogramming (Periera et al., 2020; Anderson and Perona, 2014; Hayden et al., 2021). By ethogramming, we mean the classification of pose sequences into specific behavior into ethologically meaningful categories such as walking, foraging, grooming, and sleeping. Currently, constructing an ethogram requires the delineation of ethogrammatical categories, which involves the time-consuming and careful annotation of behavior by highly trained human observers. Human-led ethogramming is slow, extremely costly, error-prone, and susceptible to characteristic biases (Anderson and Perona, 2014; Kardish et al., 2014; Holman et al., 2015; Tuyttens et al., 2014). For these reasons, it is simply impractical for even moderately large datasets, collected either in an open environment or the home cage (Womelsdorf et al., 2021). These kinds of datasets require automated alternatives. Automated ethogramming requires both high quality behavioral tracking and novel methods applied to tracked data that result in detection of meaningful categories. Such techniques have not, until recently, existed for primates (Hayden et al., 2021). Our methods take the raw information needed for ethogramming - pose data - and infer actions and higher-level categories from it. As such, they provide the first step towards automated ethogramming in primates. There are a number of steps that may be taken in the future to increase and refine the repertoire of discovered behaviors, including improvements in image capture resolution, more accurate and robust models of pose, and more varied experimental designs that induce more varied behaviors. We are particularly optimistic about the potential benefits of ethogramming for systems neuroscience. Relating behavior to neural circuits and networks is an important goal in the field, so being able to quantify behavior more rigorously– without sacrificing freedom of movement or naturalness is likely to be invaluable for future studies.

## METHODS

### Animal procedures

Animal procedures were designed and conducted in compliance with the Public Health Service’s Guide for the Care and Use of Animals and approved by the institutional animal care and use committee (IACUC) of the University of Minnesota. Two male rhesus macaques (*Macaca mulatta*) served as subjects. Animals were habituated to laboratory conditions, trained to enter and exit the open arena, and then trained to operate the water dispensers. We placed a cranial form-fitted Gray Matter (Gray Matter Research) recording chamber and a 128-channel microdrive recording system (SpikeGadgets) over the area of interest. We verified positioning by reconciling preoperative magnetic resonance imaging (MRI) as well as naive skull computed tomography images (CT) with postoperative CTs. Animals received appropriate analgesics and antibiotics after all procedures. The planning of the chamber and subsequent image alignment was performed in 3D slicer. Brain area segmentation followed the macaque D99 parcellation in NMT space (Saleem et al., 2021).

### Electrophysiology

Recordings were made with a 128-channel microdrive system (SpikeGadgets), targeting a wide swath of the prefrontal cortex ranging from OFC to PMd. Each electrode was independently moveable along the depth dimension. Neural recordings were acquired with a wireless datalogger (HH128; SpikeGadgets). The datalogger was triggered to start recording with a wireless RF transceiver (Spikegadgets), and periodically received synchronization pulses. Data were recorded at 30 kHz, stored on a memory card for the duration of the experiment, and then offloaded after completion of the session. Each feeder had local code running the experiment.

Task events triggered a TTL pulse, as well as a wireless event code. A dedicated PC running custom code controlled all feeders, and aggregated event codes. Syncing of all data sources as accomplished via the Main Control Unit (MCU; Spikegadgets), which received dedicated inputs from the pose acquisition system (see below), and feeders. Recording sessions were initiated and controlled by Trodes software (Spikegadgets). After neural recordings were offloaded, they were synced with other sources of data via the DataLogger GUI (Spikegadgets).

Recordings were performed for 4-6 days weekly for a period of 4-6 months. For an initial period of 2-4 weeks, we lowered up to 10 electrodes in each session until each had punctured the dura and their position was well-within cortex as seen from the fMRI reconstruction. Subjects still performed experiments during this time, but as the signal was noisy, no recordings were performed during this time.

A typical recording day consisted of multiple stages, including electrode adjustment, an experimental session, and extraction of the recorded signal. For the duration of the experiment, on each day, we tracked yields on each electrode and visually assessed the quality of the signal. If an electrode had poor yields for up to 5 days in a row, we would lower it up to 1 mm (or more if it was intended to move to a new area).

Spike sorting was performed in a semi-supervised fashion with a modified version of the wave_clust software (Chaure et al 2018). After an automatic phase which extracted putative units, lab members then curated the output. Units were retained on the basis of waveform shape, inter-spike interval distribution, and temporal stability.

Subdivisions of the brain were collapsed to anatomical areas, listed below as defined in the D99 parcellation of the NMT atlas (Saleem et al., 2021):

- dACC: 24a’, 24a, 24b, 24b’, 24c, 24c’
- vlPFC: 45a, 45b, 46d, 46v, 46f, 12r
- dlPFC: 8ad, 8bd, 8av, 8bs, 9d, 8bm, 9m
- SMA: F3, F6
- PMd: F1, F2, F5, F7, F4
- OFC: 13b, 13m, 13l, 12l, 12m, 12o, 11l, 11m

### Task

There were four water dispensing stations (“patches”) available with programmed delivery schedules. Each patch delivered a fixed amount of 1.5 mL per lever press. The first four presses, regardless of patch sequencing, were rewarded with water delivery. The fifth lever press was unrewarded and led to a 3-minute deactivation of the patch. That is, animals could press fewer than five times, leave to engage with a different patch, and return to the same patch in the state they left it. No reset or deactivation was applied if the animal left the patch. A patch was only reset if the subject pressed the lever a fifth time and waited 3 minutes for it to reactivate. Each rewarded press followed the same programmed sequence.

An activated patch was indicated by a fully blue display. A lever press changed the display to white with a green plus-sign in the center, an auditory cue was played, and a solenoid opened to dispense reward. After dispensing the solenoid closed, the auditory cue ended and the green plus-sign disappeared. The screen remained white for two additional seconds before the screen turned blue again to indicate the availability of another lever press. The fifth lever press was instead followed by the screen immediately turning white, with no visual or auditory reward cue and no water delivery. However, it is important to note that task engagement was not required; a subject could choose not to engage with the patches for the entirety of the session.

Otherwise, the measured behavior was simply the free movement of the subject through the arena. On average, subjects pressed levers 136 +/-8 times (subject Y: 166 +/-7, subject W: 107 +/-9), amounting to roughly 204 mL of water, per session, received for interacting with patches. Prior and concurrent chaired task training of these subjects included two risky choice tasks (Azab and Hayden, 2018; Farashahi et al., 2019; Yoo et al., 2020) and a simpler choice task (Blanchard et al., 2014; Ebitz et al., 2019).

### Pose acquisition

Images were captured with 62 cameras (Blackfly, FLIR), synchronized via a high-precision pulse generator (Aligent 33120A) at a rate of 30 Hz. The cameras were positioned to ensure coverage of the entire arena, and specifically, so that at least 10 cameras captured the subject with high-enough resolution for subsequent pose reconstruction, regardless of the subject’s position and pose. Images were streamed to one of 6 dedicated Linux machines. The entire system produced about six TB of data for a two hour session. After data acquisition, the data were copied to an external drive for processing on a dedicated Linux server (Lambda Labs).

To calibrate the camera’s geometries for pose reconstruction, a standard recording session began with a camera calibration procedure. A facade of complex and non-repeating visual patterns (mixed art works and comic strips) was wrapped around two columns of barrels placed at the center of the room, and images of this calibration scene were taken from all 62 cameras. These images were used to calibrate the camera geometry (see below). This setup was then taken down, and the experiment began.

### Pose reconstruction

We first extracted parameters relating to the cameras’ geometry for the session. To this end, we used a standard structure-from-motion algorithm (*colmap;* Schonberger and Frahm, 2016) to reconstruct the space containing the 3D calibration object and 62 cameras from the calibration images, as well as determine intrinsic and extrinsic camera parameters. We first prepared images by subtracting the background from each image in order to isolate the subject’s body. Then, 3D center-of-mass trajectories were determined via random sample consensus (RANSAC). Finally, the 3D movement and subtracted images were used to select and generate a set of maximally informative cropped images, such that the subject’s entire body was encompassed. To reduce the chance that the tire swing would bias pose estimation, we defined a mask of pixels to ignore that encompassed the tire’s swinging radius.

Next, we inferred 3D joint positions using a trained convolutional pose machine (CPM; Bala et al., 2020). We used a loss function that incorporated physical constraints (such as preserving limb length, and temporal smoothness) to refine joint localization. We found residual variability in limb length across subjects after reconstruction, between subjects, particularly for the arm, resulting in poses that were highly specific to individual subjects. To prevent subject-specific limb lengths from biasing subsequent behavior identification, we augmented the original 13 inferred landmarks to include two new ones (positions of left and right elbows) using a supplementary trained CPM model (method described in Bala et al., 2020). Thus, the augmented reconstruction resulted in 15 annotated landmarks for each image.

### Pose preprocessing

We applied a number of smoothing and transformation steps to the 3D pose data. First, we transformed the reconstructed space to a reference space that was measured using the Optitrack system (Bala et al., 2020). Then, we ignored any frame where a limb was outside the bounds of the cage due to poor reconstruction, or residual frames where subject poses were still subject to collapse (defined as where the mean limb length < 10 cm). Next, we interpolated over any segments of missing data (lasting at most 10 frames, or 0.33 sec) using a piecewise cubic interpolation. After this, we also removed any segments lasting less than 30 seconds. Note that only a small number of frames were removed after this whole procedure; specifically, 0.64% of frames on average were ignored. Finally, we rescaled any limbs in frames where the limb length was >3 standard deviations above the session mean to be at most 3 SDs long. Finally, data was smoothed with a Hampel (median) filter over 5 samples.

### Feature engineering

To discover behaviors, we next extracted a number of informative kinematic features, calculated over short time windows. Relevant parameters were calculated for one session from subject Y that exhibited a wide range of behaviors (similar results were obtained using other sessions), and then applied to every other session from both subjects.

Some features were in the *world-reference frame*. Each of these features, unless otherwise noted, was smoothed with gaussian of length 10, 30, and 60 frames (0.3, 1, and 2 sec.), resulting in a total of 8×3=24 features.

- *<u>Speed</u>*: Calculated as the absolute of the numerical derivative of the center-of-mass (COM; defined as the midpoint between the hip and neck joint) of the subject.
- *<u>Ground Speed</u>*: COM speed but just along the ground plane
- *<u>Height Speed</u>*: COM speed but just along the gravity dimension
- Height Velocity: Numerical derivative of COM along the gravity dimension
- *<u>Perpendicularity</u>*: scalar representing how vertically oriented the animal was. Calculated as the norm of the cross product of the spine (hip to neck vector) and gravity dimension. A value of 0 means the subject was vertically oriented, while 1 represents horizontal orientation.
- *<u>Height</u>*
- *<u>Limb-speed variability (height)</u>*: helps to differentiate ballistic from non-ballistic movements (e.g jumping). Obtained by calculating the speed of major joints (right hand, left hand, right foot, left foot, hip, neck), and then, for each frame, the standard deviation among these.
- *<u>Limb-speed variability (ground)</u>*: same as above, but calculating speed along the ground plane.

Next, we calculated a set of features in the *egocentric reference frame*. To this end, we normalized the orientation of poses on individual frames in a two step procedure; first, by aligning poses to face the same direction, and second, by adding back rotation corresponding to the perpendicularity of the spine relative to gravity. First, we transformed each pose to face a common direction. To do this rotation, we first defined two vectors, one corresponding to the spine (neck to hip landmarks), and the other to the expanse of the shoulders (left and right shoulder landmarks, which was then centered on the neck landmark). Poses were then rotated such that the plane defined by these vectors faced the same direction (in essence, so that the torso faced the same direction). Next, we rotated poses such that the spine had the same original orientation; in other words, if the first step aligned poses where the spine was either perpendicular or parallel to face the same direction, this step undid that rotation. We found the angle of rotation by comparing the spine (hip to neck) vector with that of gravity, and then rotated poses along the sagittal plane (ie splitting the subjects body into left and right). After this procedure, poses were aligned to the same direction, but with their original orientation of the spine (e.g if sitting or walking, poses would face in the same direction, but in the first case, the spine is vertical, while in the second, it is horizontal). From these egocentric poses, we calculated the following 67 features:

- *<u>Joints</u>*: we calculated the major sources of variation of the major joints (right hand, left hand, right foot, left foot, hip, neck). We performed a principal component analysis (PCA) of the xyz coordinates of each of these joints, and extracted the top 15 PCs.
- *<u>Joint periodicity</u>*: to ascertain periods of periodicity in poses, we obtained a time-frequency decomposition of each of the joint PCs. We first whitened the time series by performing a smooth differentiation using a Savitsky-Golay filter of order 3 and length 5 (matlab function: *sgolay*). Then, we obtained the time-frequency decomposition using Morlet wavelets via the Fieldtrip toolbox (https://www.fieldtriptoolbox.org/) function *ft_freqanalysis*, 5 cycles and 3 standard deviations of the gaussian. We obtained 100 frequencies, equally and logarithmically spaced between 0.1 and 15 Hz. We then obtained power values, and clipped them at a maximum of the 99.9th percentile the entire session. Finally, we reduced the dimensionality of this representation using PCA with 20 components.
- *<u>Segment Length</u>*: Segments were defined thus: right hand to right shoulder, left hand to left shoulder, hip to right foot, hip to left foot, neck to hip, right hand to left hand, left hand to left foot, left foot to right foot, and right foot to right hand. Segment lengths were smoothed with gaussian windows of length 10, 30, and 60 frames.
- *<u>Segment Length Periodicity</u>*: We then obtained the time-frequency representation of the (unsmoothed) segment lengths. Spectral decomposition and dimensionality reduction was performed as described above.

This process resulted in a 91-dimensional time-resolved feature set of pose kinematics. To avoid imbalance effects in embedding due to differently scaled features, we normalized each set of features described above. For each whole set of features, we calculated a robust z-score (using the median and median absolute deviation, instead of the mean and standard deviation).

### Action identification via embedding and clustering

We created behavioral maps by embedding the extracted kinematic features into two dimensions. As a first step, for computational efficiency, we first constructed a training set using data from subject Y. Specifically, for each session in subject Y, we constructed the training set by sampling every 6 points (i.e. every 200 ms). We oversampled rare events, either where the instantaneous speed was > 2, or the height of the subject’s was > 3. Roughly speaking, these corresponded to ballistic movements (such as jumping), and climbing.

We then found a 2-dimensional embedding of the training set using Uniform Manifold Approximation and Projection (UMAP; McInnes et al., 2014). We used a euclidean distance metric, and set parameters *min_dist=0*.*1, n_neighbors=200, and set_op_mix_ratio=0*.*25*, which we found to be a good balance between separating dissimilar behaviors, while combining similar ones.

To define behavioral clusters, we first estimated the probability density at 200 equally interspersed points both in the first and second UMAP dimensions. This produced a smoothed map of the pose embeddings, with clearly visible peaks (**Figure 2B**). We then employed the *watershed algorithm* on the inverse of this smoothed map (Berman et al., 2014). This algorithm defines borders between separate valleys in the (inverse of) the embedding space. Thus, the algorithm determines sections of the embedding space with clearly delineated boundaries (i.e. clusters). Samples were then assigned a cluster label according to where they fell within these borders. Using this procedure, we found a total of 59 behavioral clusters.

Finally, we re-embedded and assigned cluster labels to every sample in both subjects. We performed a k-nearest neighbors (KNN), finding the 20 nearest neighbors (again using Euclidean distance) of each sample in both subjects to that of the training set. The position of each sample in the embedding space was the median of the 20 nearest neighbors in the training samples. On the basis of the position in UMAP space, we assigned each sample a cluster label. The KNN search was performed using the *faiss* library with GPU acceleration (Facebook; github.com/facebookresearch/faiss),.

### Assessing behavioral modularity and hierarchy

To discover how postures are organized, we employed a hierarchical clustering algorithm named *Paris* (Bonald et al., 2018), using the sknetwork library (scikitnetwork.readthedocs.io/). This algorithm employs a distance metric based on the probability of sampling node pairs and performs agglomerative clustering. Paris requires no user-defined parameters (as opposed to another popular graph clustering algorithm, Louvain, which can perform hierarchical clustering according to a user-supplied resolution parameter). It is equivalent to a multi-resolution version of the Louvain algorithm (Bonald et al., 2018). The result of this algorithm is a dendrogram describing the relation between different action transitions (which we will refer to as the behavioral dendrogram). To segment action transitions into modules, we determined the modularity score (see below) for different cuts of each dendrogram for n=2 to 58 clusters). We then determined module assignment by cutting the behavioral dendrogram where the modularity score was maximized

We leveraged two important graph-theoretic metrics to assess behavioral composition:

- *Modularity Score*: The modularity score describes the degree to which postures transition within, rather than between, modules. Transition probability matrices with high modularity scores exhibit a high probability of transitions within modules, but not between modules. Modularity was calculated with the matlab function “modularity.m”.
- *Dasgputa Score*: To assess whether the graph defined by posture transitions truly reflected hierarchical organization, we calculated the Dasgputa Score (Dasgupta, 2016). The Dasgupta Score is a normalized version of the Dasgupta Cost, which defines the cost of constructing a particular dendrogram, given distances between nodes. The Dasgupta Score thus provides quantification of the quality of the hierarchical clustering. We calculated this score using the function “dasgupta_score” in the *sknetwork* library.

### Matching neural and pose data

To assess how neural activity co-varied with pose, we computed the spike-density function (SDF) of each neuron and downsampled the activity to match the pose time-series. Specifically, for each neuron, we computed the SDF using the *ft_spikedensity* function in the Fieldtrip toolbox, using a gaussian window of [-100 100] ms, sampled at a rate of 30 Hz.

As the environment allowed for free movement in the presence of a task, we sought to isolate neural activity related to postures. To this end, we regressed out neural activity related to task events at each feeder (S*creen On, Lever Press, Reward On, Reward Cue On*, and *Timeout On*), as well as the XYZ coordinates of the subject (determined as the centre-of-mass of the animals, namely the middle point between the hip and neck landmarks). Each regressor was normalized to the range [0 1]. The regressor time series was constructed at a sampling rate of 30 Hz, and smoothed with a boxcar filter of [-25 to +25 ms]. We then fit a Poisson generalized linear model (GLM) with a log link function. This model was used to predict the SDF, and the predicted SDF was subtracted from the observed SDF to obtain a residualized SDF. We excluded any cells from analysis where the Deviance was negative, due to poor model fits. Finally, for missing pose frames, activity was deleted and not analyzed. All neural analyses, unless otherwise mentioned, were performed on this residualized activity.

### Action and Action Module Encoding

To assess action encoding, we compared the average (residualized) activity of neurons, factorized by action. First, for each action segment in a session, we found averaged activity in that segment. Then, we performed a Kruskal Wallis test, with average segment activity factorized by action type. We extracted the resultant X^2^ (chi-squared) value as a metric of the encoding strength, i.e. the degree to which different actions can be separated from one another according to neuronal activity. We opted for the (non-parametric) Kruskal Wallis test as activity was highly non-normal. We assessed inter-areal differences in encoding strength using an Anova, with area as a factor.

To assess whether individual areas showed significant action encoding, we opted for a randomization test. For each unit, we first shifted the action-series of segment activities by a random amount (at least 25%, and at most 75%, of the length of the action series), and then recalculated the encoding strength. We performed this 20 times. To determine significance in each area individually, we compared the (observed) mean areal encoding strength to that of the randomized distribution.

To assess encoding for action modules, we performed the same series of analyses as outlined above but for action modules. First, in each session, we found n=2-58 action modules, as determined by different cuts of the transition probability clustering. We then remapped the action series labels according to which action module they were a part of. We then used these action module labels to find the encoding strength of each neuron.

### Assessing per-switch change in firing rate

To ascertain if neural activity was related to action switches, we computed the average activity around each switch. For each action switch in a session, we extracted neural activity in a [-1 1] segment centered on the action switch. To avoid times with multiple rapid shifts, we only considered segments where there were no other action switches in a [-0.2 0.2] sec window around the current action switch. Then, for each segment, we found the segment-normalized mean. Specifically, for each segment, we found the median and median absolute deviation of activity in the pre-switch period, and used these values to z-score normalize the entire segment.

Then, we averaged this normalized activity to obtain the per-switch segment-normalized activity of each neuron. Finally, for visualization purposes (**Figure 6A**), we averaged this activity for all units according to area. To determine if and how unit activity changed around the switch, we computed a *switching index*. Specifically, for each neuron, we obtained the average segment-normalized activity in the 1 second before and after the switch. The index was defined as the difference between the post and pre activity. To determine if switching activity differed between areas, we performed a Kruskal Wallis test.

We also considered two possible confounds that may drive our results. First, there was a large imbalance in possible action pairs. As such, per-switch differences we observed may be driven by only the most common action switches. Second, although we took care to regress out activity related to task events, some influence may nevertheless have been retained to lagged responses to task events, or poor model fits. Thus, differences in peri-switch activity may have arisen due to encoding of task events (e.g. of reward). We controlled for these possibilities in two ways. First, to address imbalances in action pairs, we computed a weighted segment-normalized activity. Specifically, we segment-normalized activity for every possible pre-switch action, and then averaged across these to obtain the *weighted segment normalized activity*. Second, to address possible residual encoding of task events, we only considered segments where subjects were not engaged with the feeders. To this end, we extracted a time-series of lever presses, and smoothed this using a 1 second boxcar filter. This gave us a time-series of *engagement probability*. Then, we selected segments where engagement probability was strictly zero. This ensured (because of the filtering step) that we only selected periods of time separated from task engagement by at least 1 sec. Finally, we then performed the same series of analyses on these *controlled segment-normalized activities*.

### Assessing the hierarchy of encoding across cortex

Our analyses suggested a gradient of postural encoding, with motor areas exhibiting more specific postural tuning, and more anterior prefrontal areas exhibiting more global encoding of the embedding space. To make concrete this observation, we performed a linear regression, predicting encoding strength corrected KL values as a function of the neurons’ location.

Specifically, the X (anterior-posterior) and Y(lateral-medial) coordinate was defined as the position of the electrode on the electrode grid, while the Z (dorsal-ventral) position was defined as the total turning depth. This was justified as electrodes were equally spaced in a grid.

To account for possible effects driven by imbalances in recording location, we performed the same analysis after binning encoding value. Specifically, we binned encoding values into a 3-dimensional grid, using 5, 10, 15, or 20 bins along each spatial dimension. We then performed the same regression, predicted (bined) encoding strength from the XYZ bin index.

We performed the same series of analyses on the *switching index* values as well, to ascertain whether changes in firing rate as a result of action switches differed according to recording location.

## Acknowledgements

We thank the Hayden/Zimmermann lab for valuable discussions. This work was supported by NIH grants R01 MH128177 (to JZ), P30 DA048742 (JZ, BH), R01 MH125377 (BH), NSF 2024581 (JZ, BH) and a UMN AIRP award (JZ, BH) from the Digital Technologies Initiative (JZ, BH), from the Minnesota Institute of Robotics (JZ).

